# *Gclc* deletion in surface-ectoderm tissues induces microphthalmia

**DOI:** 10.1101/700591

**Authors:** Brian Thompson, Ying Chen, Julien Philippe, David Anderson, Jaya Prakash Golla, Emily Davidson, Nicholas Apostolopoulos, Kevin Schey, Nicholas Katsanis, David J. Orlicky, David Thompson, Vasilis Vasiliou

## Abstract

Glutamate cysteine ligase catalytic subunit (Gclc) is the catalytic subunit for the glutamate-cysteine ligase (Gcl) enzyme. Gcl catalyzes the rate limiting step in glutathione (GSH) synthesis. *Gclc* is highly expressed in the developing eye. To define the regulatory role of *Gclc* in eye development, we developed a novel, *Le-Cre* transgene-driven, *Gclc* knockout mouse model. *Gclc*^*f/f*^/*Le-Cre*^*Tg/−*^ mice present with deformation of the retina, cornea, iris, and lens, consistent with a microphthalmia phenotype. Controlling for the microphthalmia phenotype of *Gclc*^*wt/wt*^/*Le-Cre*^*Tg/−*^ mice revealed that *Gclc*^*f/f*^/*Le-Cre*^*Tg/−*^ mice have a more severe microphthalmia phenotype. Thus, the loss of *Gclc* expression exacerbates the microphthalmia phenotype in *Le-Cre* mice. *Gclc*^*f/f*^/*Le-Cre*^*Tg/−*^ eyes present with reduced retinal and lens epithelium proliferation and increased lens cell death. Imaging mass spectrometry of ocular tissues revealed changes in the intensity and distribution of several lipid species and proteins in the retina and corneas of *Gclc*^*f/f*^/*Le-Cre*^*Tg/−*^ eyes. Lastly, using splice-blocking morpholinos and CRISPR/Cas9, we created two *gclc* knockdown zebrafish models, both of which display a microphthalmia phenotype. Combined, the mouse and zebrafish results indicate that, in chordates, *Gclc* has a conserved role in regulating eye development. In summary, these novel animal models are useful tools for elucidating the mechanisms involved in microphthalmia development.

## Introduction

Microphthalmia is a debilitating congenital malformation that affects around 1 in 7000 births [1–3]. In individuals with microphthalmia, one (unilateral) or both (bilateral) eyes are abnormally small (e.g., in adults, a corneal diameter less than 10 mm and an antero-posterior diameter of the globe less than 20 mm [4]). Microphthalmia can be classified into two main categories: total microphthalmia or partial microphthalmia [5]. In total microphthalmia, both the anterior and posterior segments are shortened. In contrast, only either the anterior *or* the posterior segment are abnormally small in partial microphthalmia. In partial microphthalmia cases where the anterior segment is abnormally small, lens development is often impaired. Lens development is regulated by a myriad of extracellular signaling pathways and gene regulatory networks (reviewed here [6]). Approximately 80% of microphthalmia cases can be explained by pathogenic mutations in *SOX2*, *OTX2*, *PAX6*, *VSX2*, *RAX*, *FOXE3*, *STRA6*, *ALDH1A3*, or *RARB* [3, 7]. The remaining 20% of microphthalmia cases may be explained by exposure to environmental factors, such as pharmaceuticals (e.g, thalidomide [8]), ionizing radiation exposure [9, 10], fungicide exposure (e.g., benomyl [11]) and gestationally-acquired infections [12], or other factors such as maternal vitamin A deficiency [13, 14]. Blindness or limited vision is observed in individuals with the severest forms of microphthalmia. Indeed, microphthalmia is a major cause of childhood blindness, accounting for ≈11% of the cases [1]. Currently, there is no cure for the severe loss of vision associated with microphthalmia. While surgical treatments exist, they only address the cosmetic abnormalities associated with microphthalmia [15]. However, recent research demonstrating the efficacy of nonsense suppression (inhibiting the effect of a nonsense mutation by pharmacologically increasing the likelihood of a near cognate aminoacyl-tRNA substitution) to rescue lens development when administered postnatally in a *Pax6*-deficient mouse model has provided hope that treatments for microphthalmia can be developed [16]. Clearly, a more complete understanding of the mechanisms regulating lens development should accelerate the discovery of preventative strategies and/or novel therapeutic treatments for microphthalmia.

Glutathione (GSH), an intracellular antioxidant, is highly abundant in the eye, with concentrations of 20mM or more occurring in the lens cortex and epithelium [17, 18]. It is important for physiological detoxification of electrophiles and oxidants in the lens cortex and epithelium [19]. Through these functions, GSH maintains lens clarity and, as such, a decline in GSH levels is associated with cataract development [20, 21]. GSH biosynthesis involves a two-step enzymatic process [22] involving glutamate-cysteine ligase (GCL) followed by glutathione synthase (GSS). GCL, the rate limiting step in GSH synthesis, comprises a catalytic subunit (GCLC) and a modifier subunit (GCLM). Impaired GCLC or GCLM function limits GSH synthesis and reduces intracellular GSH concentrations. Thus, several *Gclm* [23] and *Gclc* [24–26] -deficient mouse models have been created to explore the role of GSH in physiological and pathophysiological conditions. For example, such mouse models have identified a role for GSH in protection against steatosis and neurodegeneration [24, 27]. A Lens Glutathione Synthesis KnockOut (LEGSKO) mouse model revealed suppression of *Gclc* expression induced age-related nuclear cataract development [26]. In this model, *Gclc* deletion occurs in the lens at embryonic day (E) 10.5 [26]. However, the role of *Gclc* and GSH in the early stages of lens development (i.e., prior to E10.5) remains uncharacterized.

We have developed novel mouse and zebrafish models in which GSH synthesis is reduced early in lens development. This was accomplished by genetically ablating *Gclc* in the surface-ectoderm derived tissues (i.e., cornea, lens, conjunctiva, eyelid) by cross-breeding *Gclc* floxed mice (*Gclc*^*fl/fl*^) mice [24] with mice hemizygous for the *Le-Cre* transgene [28]. The *Le-Cre* transgene utilizes the *Pax6-P0* promoter to limit ocular *Cre* expression to only the lens, conjunctiva, and cornea starting at E9.5 [28]. In addition, since GSH is abundant in zebrafish [29] and *gclc* is highly expressed in the developing zebrafish eye [30], we postulated that it has a conserved role in regulating zebrafish eye development. To evaluate this possibility, we developed novel *gclc*-suppressed zebrafish models using splice-blocking morpholinos or CRISPR/Cas9. Herein, we present the phenotypic characterization of targeted GSH synthesis reduction in surface ectoderm-derived tissues during early mouse and fish eye development.

## Methods

### Generation of Surface-Ectoderm Specific *Gclc* Knock-out Mice

In order to delete *Gclc* from surface ectoderm-derived tissues, *Gclc* floxed (*Gclc*^*f/f*^) mice (C57BL/6J background) were bred with *Le-Cre* (FVB/N background) transgenic mice (*Le-Cre*^*Tg/−*^). The resultant *Gclc*^*f/f*^/*Le-Cre*^*Tg/−*^ (*Gclc* KO) mice have ocular *Cre* recombinase expression restricted to only the conjunctiva, cornea, eyelid and lens from embryonic day 9 (E9) [28]. The generation of the *Le-Cre (Tg(Pax6-cre,GFP)1Pgr)* transgenic and *Gclc*^*f/f*^ mice have been previously described [24, 29]. These mice [28] were originally produced by Dr. Ruth Ashery-Padan (Department of Human Molecular Genetics and Biochemistry, Sackler Faculty of Medicine, Tel Aviv University), and were obtained from Dr. David Beebe (Department of Ophthalmology and Visual Sciences, Washington University School of Medicine). *Gclc* HET were continuously crossed with *Gclc*^*f/f*^ mice to convert their genomic background towards C57BL/6J and generate the three experimental mouse genotypes; *Gclc*^*f/f*^/*Le-Cre*^−/−^ (*Gclc* WT), *Gclc*^*wt/f*^/*Le-Cre*^*Tg/−*^ (*Gclc* HET), and *Gclc*^*WT/f*^/*Le-Cre*^*Tg/−*^ (*Gclc* KO). All data presented in this paper was generated from B6/FVB mixed background mice. Mice were group-housed and maintained on 12-hour light-dark cycles, with food and water available *ad libitum.* All experiments were performed in strict accordance with the National Institutes of Health guidelines, with protocols approved by the Yale University Institutional Animal Care and Use Committee.

**Table 1.** Definitions of abbreviations used for mouse genotypes.

| Abbreviation | Genotype |
| --- | --- |
| *Gclc* WT | *Gclc*<sup>f/f</sup>/*Le-Cre*<sup>-/-</sup> |
| *Gclc* HET | *Gclc*<sup>wt/f</sup>/*Le-Cre*<sup>Tg/-</sup> |
| *Gclc* KO | *Gclc*<sup>f/f</sup>/*Le-Cre*<sup>Tg/-</sup> |
| *Gclc* WT/Cre | *Gclc*<sup>wt/wt</sup>/*Le-Cre*<sup>Tg/-</sup> |

### Generation of *Le-Cre* Transgene Control Mice

To generate *Le-Cre* transgene control mice, we bred *Gclc*^*f/f*^/*Le-Cre*^*−/−*^ (*Gclc* WT) mice with *Gclc*^*wt/f*^;*Le-Cre*^*Tg/−*^ (*Gclc* HET) mice. The F1 mice (*Gclc*^*wt/f*^/*Le-Cre*^*Tg/−*^ and *Gclc*^*wt/f*^/*Le-Cre*^*−/−*^) were then interbred to produce F2 mice of the *Gclc*^*wt/wt*^/*Le-Cre*^*Tg/−*^ (WT/Cre) genotype. These WT/Cre mice served as *Le-Cre* transgene control mice.

### Genotyping

Genomic DNA, obtained from a 2 mm ear punch collected at post-natal day (P) 14, was extracted using DirectPCR Lysis Reagent (Tail) (Viagen Biotech, Los Angeles, CA) and PCR Grade Proteinase K (Sigma-Aldrich) according to the manufacturer’s protocol. Briefly, the ear punch sample was placed in 100 μL DirectPCR Lysis Reagent (Tail) containing 1 μL PCR Grade Proteinase K. The sample was incubated overnight at 55°C, and subsequently incubated at 85°C for 45 minutes. Two separate PCR reactions were needed to properly describe each individual animal: one detected the *Gclc(wt)* and *Cre(Tg)* alleles (left lane), and the other detected the *Gclc(f)* allele (right lane). *Gclc* alleles were determined using *5’-CTATAATGTCCTGCACTGGG* and *5’-TAGTGAACGCTGTTAAAGG* and *5’-CGGGTGTTGGGTCGTTTGT* and the presence of the *Cre* transgene as identified using *5’-GCGTCTGGCAGTAAAAACTATC* and *5’-GTGAAACAGCATTGCCACTT*, as previously described *[24]*.

### Histopathological Examination and Immunohistochemical Staining

E14.5 mice were collected, genotyped, and fixed overnight in 10% neutral buffered formalin (Sigma-Aldrich). Ocular tissues were processed (i.e., dehydrated) and embedded in paraffin wax and sectioned (5 μm thickness) onto slides by Dr. Mark Petrash (University of Colorado School of Medicine). A section from both eyes of each mouse was stained with hematoxylin and eosin (H&E). Immunostaining was conducted using the TSA Plus Biotin Kit (Perkin Elmer, Waltham, MA) according to the manufacturer’s protocol. Briefly, slides were blocked with TNB Blocking Buffer (Perkin Elmer, Waltham, MA) for 30 minutes at room temperature. Subsequently, slides were incubated overnight with primary antibody in TNB at 4°C. After primary antibody exposure, the slides were washed 3 times for 5 minutes with TNT buffer. Subsequently, the slides were incubated with horseradish peroxidase (HRP) conjugated secondary antibody for 30 minutes at room temperature. Slides were then washed 3 times for 5 minutes with PBST (0.05% Tween-20, 1 X PBS) at room temperature. Next, the slides were incubated with TSA Plus working solution (Perkin Elmer, Waltham, MA) for 10 minutes at room temperature. The slides were subsequently incubated in streptavidin conjugated (SA)-HRP (diluted 1:100 in TNB) for 30 minutes at room temperature, Harris hematoxylin counterstain was applied, and mounted for imaging. Antibodies: PAX6 (Abcam, ab195045), GCLC (Abcam, ab53179), Ki67 (Abcam, ab21700), β-Catenin (Abcam, ab32572), and Rabbit IgG H&L HRP (Abcam, ab6721). All antibodies were used at a concentration of 1:200 unless otherwise stated. TUNEL staining was performed by the Yale University Pathology Tissue Services.

### Quantification of GSH and GSSG

To quantify GSH and GSSG, postnatal day (P) 21 *Gclc* KO and *Gclc* WT mice were anesthetized and eyes enucleated and immediately assayed using the GSH/GSSG Ratio Detection Assay Kit (Fluorometric-Green) (Abcam) according to the manufacturer’s protocol. In brief, the whole eye tissue was rinsed with Phosphate-buffered saline (137 mM NaCl, 2.7 mM KCl, 8 mM Na_2_HPO_4_, and 2 mM KH_2_PO_4_) resuspended in ice cold Mammalian Lysis Buffer (Abcam) and homogenized using Tissuelyser ^**TM**^ (QIAGEN, Venlo, Netherlands) at a frequency of 30 Hz for 2 minutes at 4°C. The homogenate was centrifuged at 10,000 g for 30□minutes at 4°C and GSH and GSSG levels were measured in the supernatant. To measure GSH and GSSG, a one-step fluorometric reaction of samples with respective assay buffer and probes was incubated for 30□minutes protected from light. At the end of the incubation period, fluorescence intensity was monitored at EX/EM of 490/520□nm fluorescence on a microplate reader (Spectramax M3, Molecular Devices). The data are presented as the mean and standard error of the mean. Differences were determined using two-tailed Mann-Whitney test (GraphPad Prism version 7.0a for Mac OS X, GraphPad Software, La Jolla California, USA) with p < 0.05 being considered significant

### Protein Extraction and Western Blotting

Total cell lysates were generated in RIPA buffer (1% Nonidet P40, 0.5% sodium deoxycholate, 0.1% SDS in PBS) by homogenization with a Tissuelyser ^**TM**^ (QIAGEN, Venlo, Netherlands) at a frequency of 30 Hz for 2 minutes at 4°C. Protein concentrations in the homogenate were quantified using the Pierce BCA Protein Assay Kit (Thermo Fisher Scientific) according to the manufacturer’s protocol. 10 μg of protein was resolved on 4-20% SDS-PAGE gradient gels (Bio-Rad, Hercules, CA) at 100 V and then transferred to a 0.2 μm nitrocellulose blot (Bio-Rad, Hercules, CA) with the Trans-blot Turbo Transfer System (Bio-Rad, Hercules, CA). The immunoblotting was performed as previously described ([31]) with antibodies against GCLC (Abcam; ab190685) and αA-crystallin (Santa Cruz; sc-28306).

### Determining Eye Mass

To determine the mass of eyes from *Gclc* WT, *Gclc* KO, and WT/Cre mice, eyes were enucleated at P21. Following removal of non-ocular tissues by dissection, the eyes were subsequently weighed (Mettler Toledo NewClassic MF). Data are represented as scatter plots. Each point is the average weight of both the left and right eye from one mouse. Differences were determined using ANOVA (GraphPad Prism version 7.0a for Mac OS X, GraphPad Software, La Jolla California, USA), with p ≤ 0.05 being considered significant.

### Preparing of Eyes for Imaging Mass Spectrometry (IMS)

P3 and P21 *Gclc* WT, *Gclc* HET, or *Gclc* KO mice were anesthetized and their eyes enucleated and frozen by placing them in a plastic weighing boat floated on liquid nitrogen. The frozen eyes were shipped to Vanderbilt University on dry ice where they were embedded by rapid freezing in 2.6 % carboxymethyl cellulose (Sigma Aldrich, St. Louis, MO, USA). This was accomplished by placing the frozen tissue into an aluminum weight boat (Electron Microscopy Sciences, Hatfield, PA, USA) on dry ice in a polystyrene container and slowly adding carboxymethyl cellulose with a pasture pipette until the eye was coated on all sides. Twelve µm sections were obtained in a sagittal orientation using a cryostat (CM3050S, Leica Biosystems, Buffalo Grove, IL). Triplicate adjacent sections were taken from the central region of the same eye and mounted onto a polylysine-coated Indium-Tin-Oxide (ITO) slide (Delta Technologies, Loveland, CO).

### IMS on Prepared Eye Tissue

Lipid imaging experiments were performed by sublimating the matrices 2,5-dihydroxyacetophenone (DHA) (Sigma Aldrich, St. Louis, MO, USA) 1,5-diaminonaphthalene (DAN) (Sigma Aldrich) onto the samples using an in-house sublimation apparatus [30]. Ten minutes at approximately 56 mTorr at 110 ºC for DHA provided a suitable coating for a 10-micron pixel size in positive ion mode. DAN was sublimated for 15 minutes under the same conditions for negative ion mode analysis. Lipid imaging experiments were performed using a rapifleX^TM^ MALDI Tissuetyper^TM^ (Bruker Daltonics, Billerica, MA) MALDI TOF instrument equipped with a 355 nm Smartbeam 3D laser operated at 10 kHz. Data were acquired in positive and negative ion mode with a 15 μm pixel size and 200 shots per pixel over a mass range from 300-1300 Da for negative ion and 200-1300 Da for positive ions.

Protein imaging experiments were performed on tissue sections that were prepared using serial washes for 30 seconds each in 70% ethanol, 100% ethanol Carnoy’s fluid (60% ethanol, 30% chloroform, 10% acetic acid (2 minutes), 100% H_2_O, and 100% ethanol. The tissue was air-dried before 2,5-DHA (Sigma Aldrich) matrix (10 mg/mL in a solution comprising 7mL acetone, 1.5 mL H_2_O, 0.4 mL trifluroacetic acid TFA, and 0.4 mL acetic acid) was applied by a HTX TM sprayer (HTX Technologies, Carrboro, NC). Four matrix spray passes were made over the tissue with a flow rate of 1.5 mL/min at 30°C at a track speed of 1300 cm/min and a track spacing of 2.5 mm. Data were acquired in positive ion mode using the rapifleX^TM^ MALDI Tissuetyper^TM^ with a 20 μm pixel size and 100 shots per pixel over a mass range of 1600-2000 Da.

MALDI images were generated using FlexImaging 5.0 (Bruker Daltonics, Billerica, MA) and data were normalized to the total ion current. Relative intensities were adjusted on an individual basis to improve image quality.

### Gene Suppression in Zebrafish Embryos

Splice-blocking morpholinos (sbMOs) targeting either the zebrafish (*Danio rerio*) *gclc* exon 1 and exon 5 were designed and synthesized by Gene Tools LLC (Philomath, OR). To determine the optimal sbMO dose, we injected 1 nL of increasing amounts of each MO (3 ng/uL, 6 ng/uL, and 9 ng/uL of injection cocktail) into EkxAB embryos (1-to 4-cell stage) harvested from natural mating. To determine MO efficiency, we used TRIzol (Thermo Fisher) to extract total RNA from embryos at 3 days post-fertilization (dpf) according to the manufacturer’s instructions. Resulting total RNA was reverse transcribed into cDNA using the Superscript III Reverse Transcriptase kit (Thermo Fisher) and used as template in RT-PCR reactions to amplify regions flanking MO target sites. 10 ng of the RT-PCR product were run on a 2% agarose gel to determine to the effect of the sbMOs. RT-PCR products were cloned (TOPO-TA; Invitrogen), and the plasmids purified from individual colonies was/were Sanger-sequenced according to standard protocols to identify the precise alteration of the endogenous *gclc* transcript.

### CRISPR/Cas9 Genome Editing in Zebrafish Embryos

We used CHOPCHOP [32] to identify a guide (g)RNA targeting exon3 of *gclc*. gRNAs were *in vitro*-transcribed using the GeneArt precision gRNA synthesis kit (Thermo Fisher) according to the manufacturer’s instructions. Zebrafish embryos were obtained from EKxAB embryos harvested from natural mating. One nL of injection cocktail (100 pg/nl gRNA and 200 pg/nl Cas9 protein (PNA Bio) in water) was injected into the embryos at the 1-cell stage. To determine targeting efficiency in founder (F0) mutants, we extracted genomic DNA from 2 days post-fertilization (dpf) embryos and PCR amplified the region flanking the gRNA target site. To estimate the percentage of mosaicism of *gclc* F0 mutants, PCR products were Sanger sequenced according to standard procedures. Cutting efficiency was assessed from sequencing chromatograms using TIDE (Tracking of Indels by DEcomposition) analysis [33].

### Phenotypic Analyses in Zebrafish

To study eye development, larval batches were reared at 28°C and imaged live at 3 days-post fertilization (dpf) using the Vertebrate Automated Screening Technology Bioimager (VAST; Union Biometrica) mounted on an AxioScope A1 (Zeiss) microscope using an Axiocam 503 monochromatic camera and Zen Pro 2012 software (Zeiss) as described previously [34]. We obtained lateral brightfield images of whole larvae using the VAST onboard camera and the Axiocam 503 camera. Eye area was automatically measured using FishInspector [35]. The FishInspector Framework provides algorithms and routines for automatic feature-detection from images recorded by the VAST Bioimager Platform; this allowed for unbiased evaluation of statistically-relevant sample sizes. The data are presented using scatter plots. Differences were determined using ANOVA with Tukey correction (GraphPad Prism version 8.0 for Mac OS X, GraphPad Software, La Jolla California, USA), with p < 0.05 being considered significant.

## Results

### Lens-specific *Gclc* deletion reduces whole eye GSH content

We successfully generated three genotypes, *Gclc* WT, *Gclc* HET and *Gclc* KO, by crossing *Gclc*^*f/f*^ mice with *Le-Cre*^*Tg/−*^ (Fig. 1A). As expected from previous results [24], the conditional deletion of *Gclc* did not cause embryonic lethality, i.e., all three genotypes were born in the expected Mendelian frequency (data not shown). Immunohistochemical staining of GCLC in the eyes of E14.5 mice confirmed that GCLC is abundant in the developing eye, and GCLC expression is absent from the surface ectoderm-derived tissues of *Gclc* KO mice (Fig. 1B). Western blot analysis at P0 confirmed the absence of GCLC expression in the lenses of *Gclc* KO mice. Interestingly, *Gclc* KO retina showed decreased GCLC expression (Fig. 1B and 1C). The lenses of *Gclc* Het mice have reduced GCLC expression compared with those of *Gclc* WT mice (Fig. 1C). GCLC binds to GCLM to form the GCL holoenzyme, which catalyzes the rate-limiting step in GSH synthesis. As expected, whole eye levels of GSH and GSSG were markedly reduced in *Gclc* KO mice (Fig. 1D). The GSH/GSSG ratio in whole eye tissue was not different between *Gclc* KO and *Gclc* WT mice (Fig. 1D).

**Figure 1.**
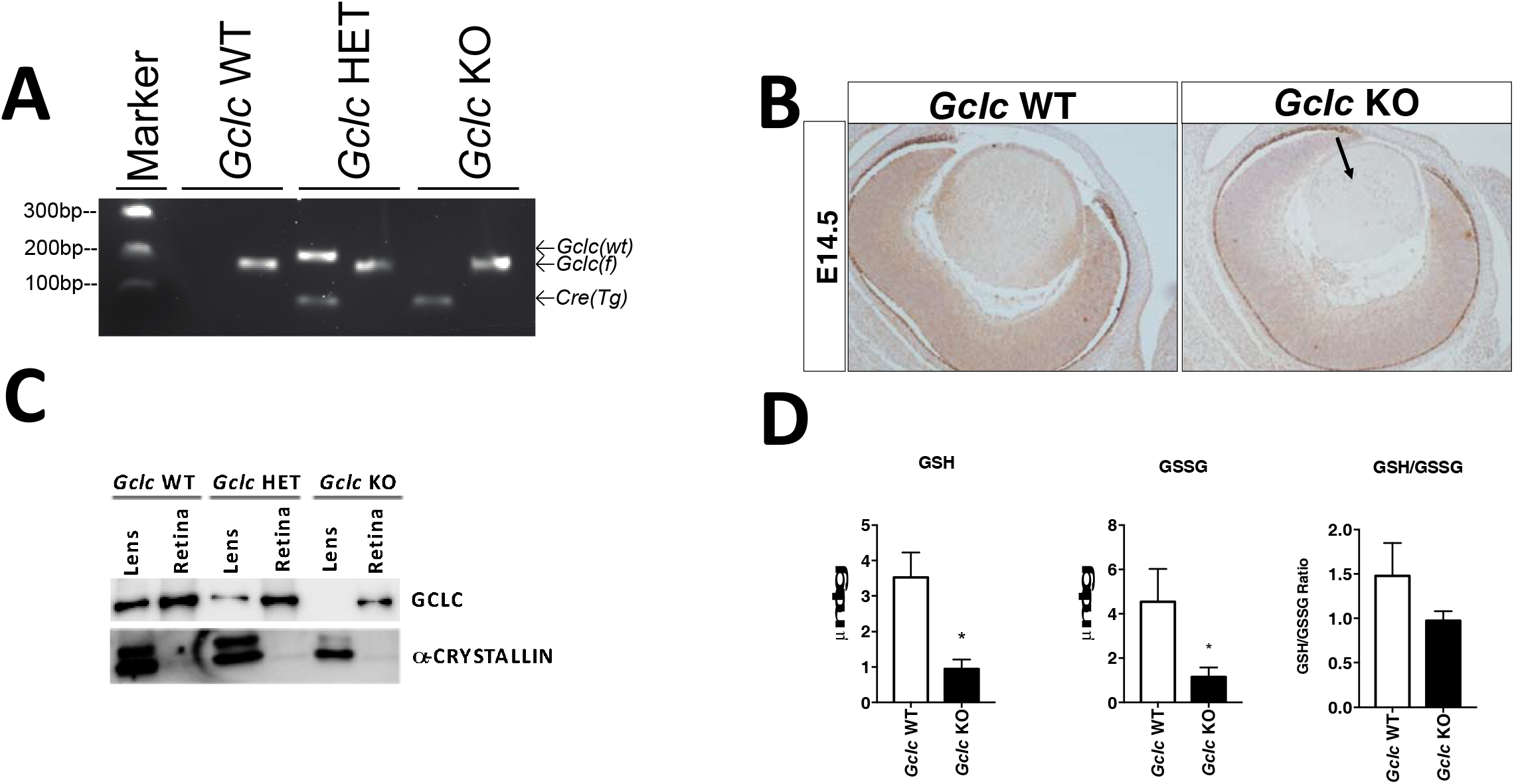
Surface-ectoderm derived tissue-specific deletion of *Gclc* reduces ocular GSH content. (**A**) Agarose gel (2%) separation of genomic DNA in P14 *Gclc*^*f/f*^/*Le-Cre*^*−/−*^ (*Gclc* WT), *Gclc*^*wt/f*^/*Le-Cre*^*Tg/−*^(*Gclc* HET), and *Gclc*^*f/f*^/*Le-Cre*^*Tg/−*^ (*Gclc* KO) mice by PCR amplification of the *Gclc wild-type (Gclc(wt))* or *Gclc floxed (Gclc (f))* alleles, or the *Le-Cre* transgene (*Cre(Tg*)). (**B**) Immunohistochemical staining of GCLC in E14.5 *Gclc* WT, *Gclc* HET and *Gclc KO* mice (Arrow marks a lens with a loss of GCLC expression; 100x magnification). (**C**) Western blot analysis of GCLC and αA-crystallin proteins in the lens and retina of P0 *Gclc* WT, *Gclc* HET and *Gclc KO* mice. The upper band observed in the blot stained for αA-crystallin represents an alternatively spliced αA-crystallin variant containing an additional exon. (**D**) GSH and GSSG levels in the whole eyes of P21 *Gclc* WT (white bars) and *Gclc* KO (black bars) mice. The ratio of GSH/GSSG is also shown. Data are presented as the mean and associated standard error of the mean from 3-4 mice. * P<0.05, two-tailed Mann-Whitney test, compared to Gclc WT.

### Suppression of *Gclc* impairs normal eye development

To characterize the morphological changes in the eyes of *Gclc* KO mice, we conducted gross morphological and histological analysis of eyes at various embryonic and postnatal stages of development (Fig. 2). *Gclc* WT eyes do not show any ocular abnormalities at any of the ages examined (Figs. 2A, D, G, J). In contrast, *Gclc* KO mice present age-dependent pathological changes in the lens, cornea, iris, and retina (Figs. 2C, F, I, L). At E14.5, *Gclc* KO mouse eyes have developed all ocular tissues; however, there is a reduction in the size of the anterior chamber (Fig. 2C, arrow) and increased cellular layers in the retina inner nuclear layer (Fig. 2C, box). By the day of birth (P1), the eyes of *Gclc* KO mice are drastically altered, with lenses that have severe fiber cell vacuolation (Fig. 2F, arrow), hyperplasia of the iris (Fig. 2F, box), and cellular hyperproliferation of the retina in the ganglion cell, inner plexiform and inner nuclear layers (Fig. 2F, star). The *Gclc* KO eye abnormalities worsen with age such that by P20, the iris has formed a thick layer that remains attached to the cornea (Fig. 2I, arrow), the lens is smaller in size and vacuolated (Fig. 2I, star), the corneas have thinner epithelial cell layers with hyperproliferation and poor differentiation (Fig. 2I, box), and retinal cellular hyperproliferation is occurring in the outer nuclear, inner nuclear, inner plexiform, and ganglion cell layers (Fig. 2I, circle). By P50, these changes are becoming more profound in *Gclc* KO mice (Fig. 2L). Interestingly, *Gclc* KO mice display bilateral microphthalmia whereas *Gclc* HET mice display only unilateral microphthalmia (note that the unilateral microphthalmia in *Gclc* HET mice is unbiased towards the right or left eye). In the microphthalmia eye only, *Gclc* HET mice display identical developmental abnormalities (Fig. 2B, E, H, K) as the *Gclc* KO mice.

**Figure 2.**
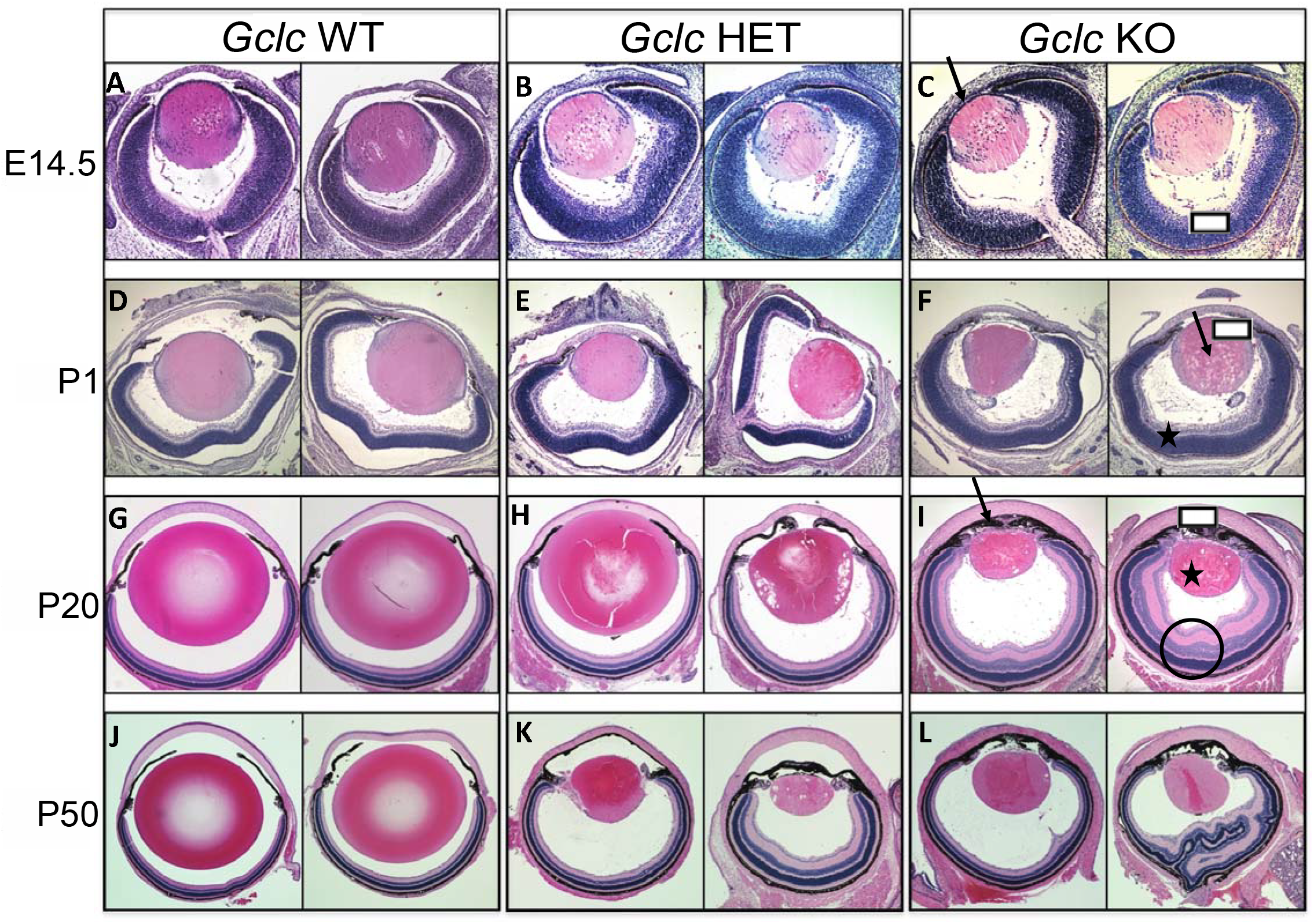
*Gclc* deletion causes severe ocular malformations. Representative images of hematoxylin and eosin-stained left (left column) and right (right column) eyes (from the same mouse at each age) from *Gclc* WT, *Gclc* HET, and Gclc KO mice at ages E14.5 (**A**-**C**), P1 (**D**-**F**), P20 (**G**-**I**), and P50 (**J**-**L**). In **C**, the arrow marks a reduction in the size of the anterior chamber and the box marks increased cellular layers of the retina inner nuclear layer. In **F**, the arrow marks vacuolated lens fiber cells, the box marks hyperplasia of the iris, and the star marks hyperproliferation of the retina ganglion cell, inner nuclear, and inner plexiform layers. In **I**, the star marks a small vacuolated lens, the box marks a cornea with abnormally thin epithelial layers with hyperproliferation and poor differentiation, and the circle marks a retina with severe hyperproliferation in the outer nuclear, inner nuclear, inner plexiform, and ganglion cell layers. Magnification = 100x.

### The microphthalmia phenotype in *Le-Cre* hemizygous mice is exacerbated by *Gclc* deletion

Our results indicate that *Gclc* appears to be required for normal eye and lens development. However, *Le-Cre* transgenic mice can display a microphthalmia phenotype, regardless of floxed alleles [36–39]. Therefore, the extent to which hemizygosity for the *Le-Cre* causes microphthalmia in our B6/FVB mixed background mice was determined (Fig. 3.) To do so, we bred *Gclc*^*f/f*^/*Le-Cre*^*−/−*^ (*Gclc* WT) and *Gclc*^*wt/f*^/*Le-Cre*^*Tg/−*^ (*Gclc* HET) mice (Fig. 1A) for at least three generations to produce *Gclc*^*wt/wt*^/*Le-Cre*^*Tg/−*^ (WT/Cre) mice, thus isolating the *Le-Cre* transgene on our B6/FVB mixed background (Fig. 3A). The combined eye weights of Wt/Cre mice and *Gclc* KO mice were less (P<0.05) than those in *Gclc* WT mice and the weights in Gclc *KO* mice were less (P<0.05) than those in WT/Cre mice (Fig. 3B).

**Figure 3.**
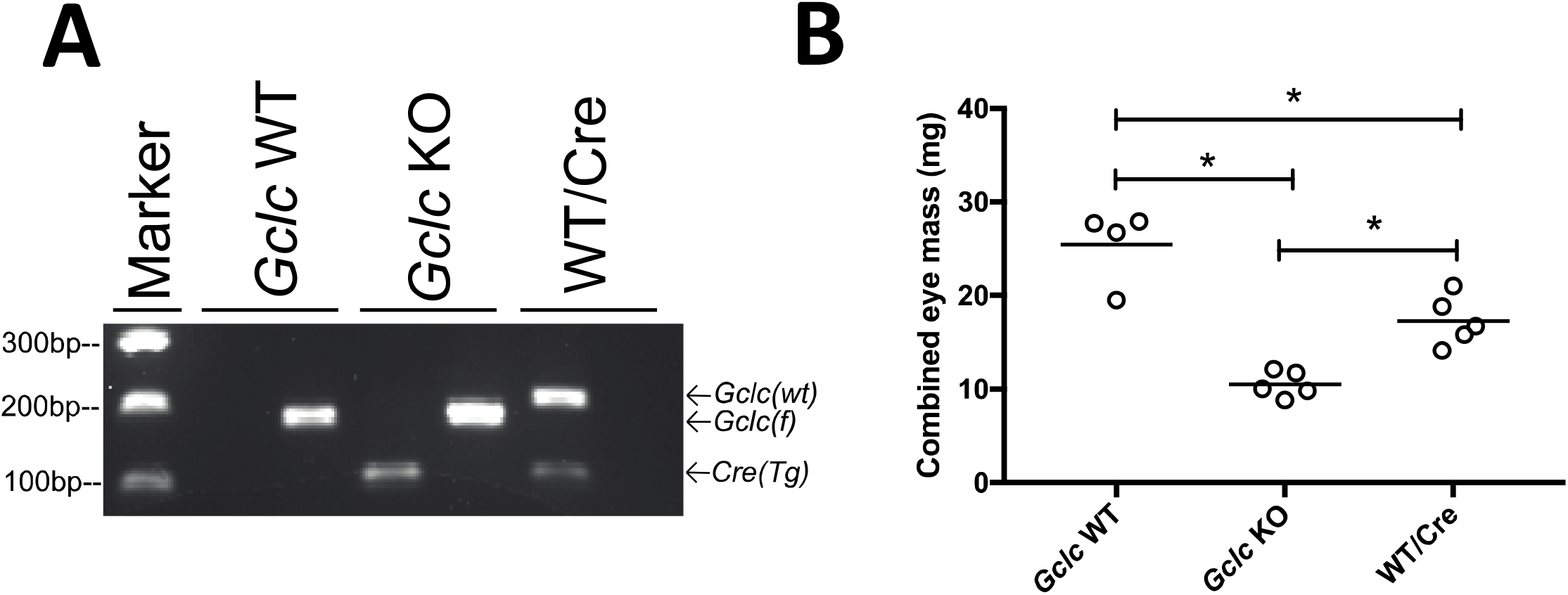
*Gclc* deletion potentiates the microphthalmia phenotype of the *Le-Cre* transgene. (**A**) Agarose gel (2%) separation of genomic DNA in P14 *Gclc*^*f/f*^/*Le-Cre*^*−/−*^(*Gclc* WT), *Gclc*^*f/f*^/*Le-Cre*^*Tg/−*^(*Gclc* HET), and *Gclc*^*wt/wt*^/*Le-Cre*^*Tg/−*^(WT/Cre) mice by amplification of the *Gclc* wild-type (*Gclc(wt)*) or *Gclc* floxed (*Gclc(f)*) alleles, or the *Le-Cre* transgene (*Cre(Tg)*). Two separate PCR reactions are needed to properly describe each individual animal: one detected the *(wt)* and *Cre(Tg)* alleles (left lane) and the other the *Gclc(f)* allele (right lane). (**B**) Combined (left and right) eye weights from P21 *Gclc* WT, *Gclc* KO and WT/*Cre* mice. Each point represents results from an individual mouse. Horizontal lines indicate the mean. * P<0.05, ANOVA, with Tukey correction, compared to other group indicated.

### *Gclc* deletion induces molecular changes within the developing eye

Immunohistochemical analysis of the eyes of E14.5 day old mice revealed molecular changes in *Gclc* KO eyes (Fig. 4). Cellular expression of Ki67 (a proliferation marker) was high throughout both the retina and lens epithelium of *Gclc* WT mice (Fig. 4A). Less staining occurred in the retina and lens epithelium (from the germinative zone to the equator) of *Gclc* KO mice (Fig. 4E). Apoptotic cells (as revealed by TUNEL staining) were observed within the germinative zone and differentiating secondary fiber cells of *Gclc* KO eyes (Fig. 4F). No differences in ocular β-catenin expression were observed between *Gclc* KO and *Gclc* WT mice (Fig. 4C & G). Lenticular Pax6 expression and distribution in Gclc KO eyes (Fig. 4H) appeared to be comparable to that in *Gclc* WT eyes (Fig. 4D). It should be noted, however, that retinal Pax6 expression was greatly reduced in *Gclc* KO eyes (Fig. 4H) relative to *Gclc* WT eyes (Fig. 4D).

**Figure 4.**
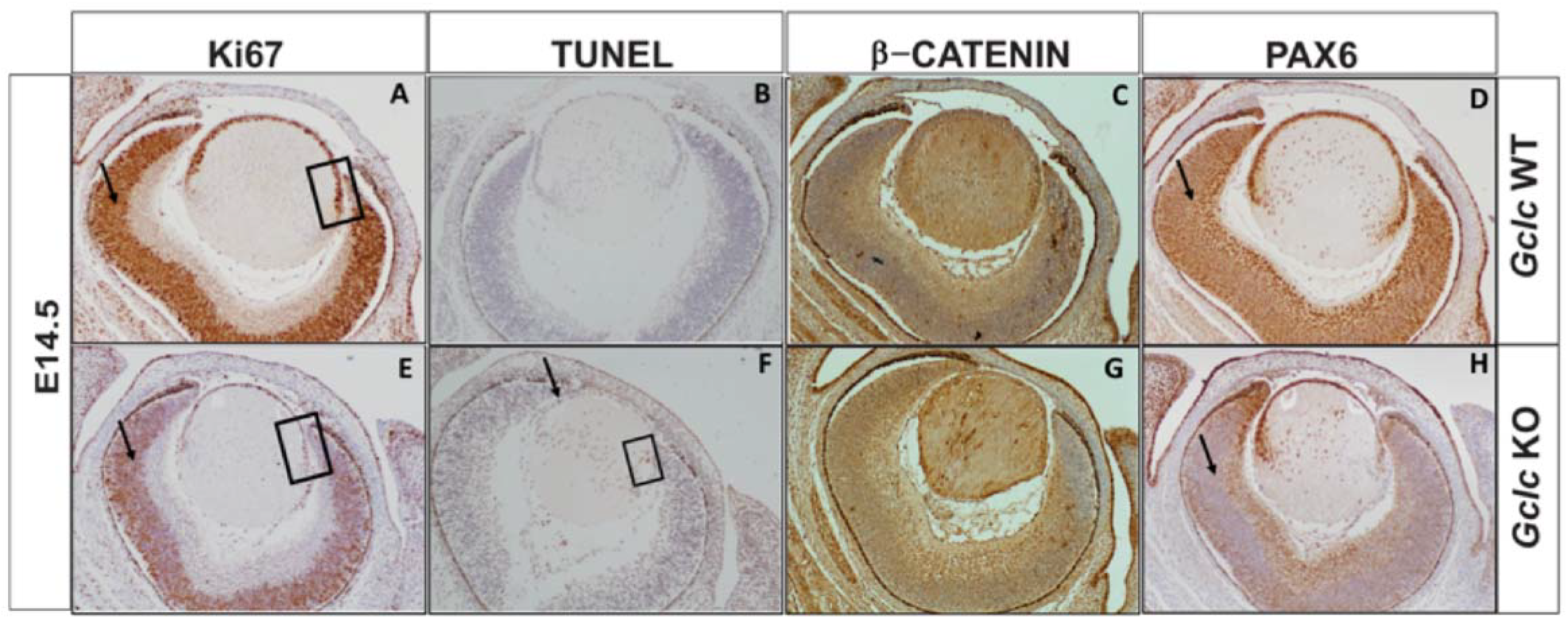
*Gclc* deletion induces molecular changes during embryonic eye development. Staining of sagittal sections of eyes from E14.5 day old *Gclc* WT (**A**-**D**) and *Gclc* KO (**E**-**H**) mice for Ki67 (**A**,**E**), TUNEL (**B**,**F**), β-catenin (**C**,**G**), or Pax6 (**D**,**H**). In **A** and **E**, the brown-staining Ki67-positive (proliferative) cells are identified in the retina (arrow) and in the lens epithelial cells (box). In **F**, TUNEL-positive (apoptotic) cells in the lens epithelium (arrow) and secondary lens fiber cells (box) are shown. In **D** and **H**, Pax6-positive cells in the retina are identified by an arrow. (Magnification = 100x)

### IMS analysis of the whole eye

Matrix-assisted laser desorption/ionization imaging mass spectrometry (IMS) was employed, in a proof-of-concept manner, to demonstrate that molecular changes in lipids and proteins can be identified within the eye of a microphthalmia mouse model (Fig. 5). Hematoxylin & eosin staining of the eyes prepared for IMS (Fig. 5A) displayed the same malformations noted previously (Fig. 2), i.e., *Gclc* KO eyes have smaller lenses that were vacuolated, exhibited cellular hyperproliferation of the retina, and a decrease in the size of the anterior chamber (Fig. 5A). Analysis of eyes from mice age P21 for changes in protein relative abundance (based on signal intensity) and spatial distribution revealed differences between *Gclc* KO and *Gclc* HET eyes relative to *Gclc* WT eyes (Fig. 5B). Specifically, thymosin β4 (at m/z 4967) signal intensity was reduced in both *Gclc* KO and HET corneal tissues in a manner that appeared to be *Gclc* gene dose-dependent, i.e., thymosin β4 was reduced to a greater extent in *Gclc* KO eyes than *Gclc* HET eyes. (Fig. 5B). Cytochrome C oxidase 7C (at m/z 5449) and histone H2B (at m/z 1378) intensities and distribution were in *Gclc* KO and *Gclc* HET mice were similar to *Gclc* WT mice (Fig. 5B); these proteins clearly illustrate the retinal malformations in *Gclc* KO eyes.

**Figure 5.**
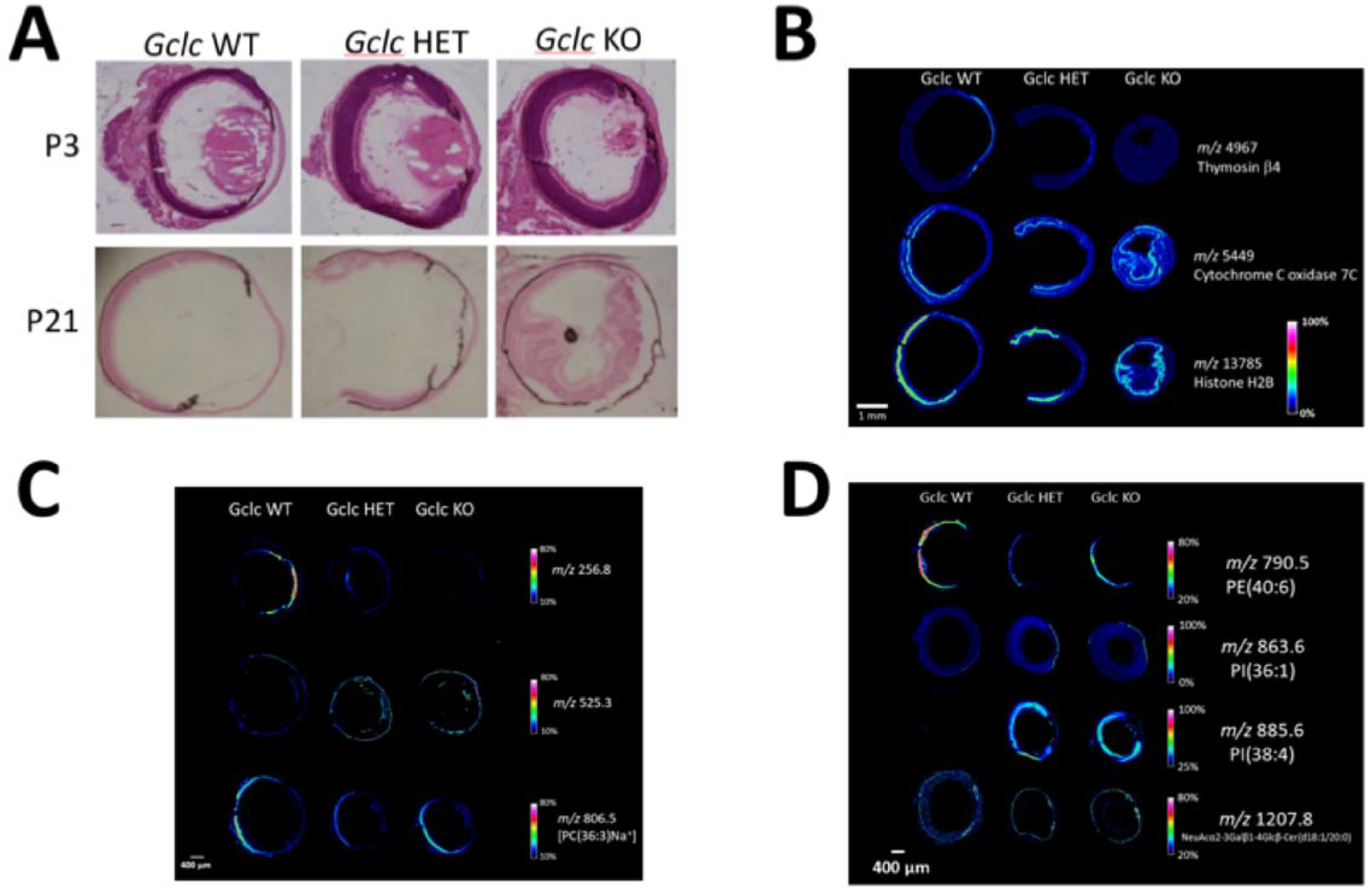
Imaging mass spectrometry (IMS) analysis of the eye. Histology and IMS were conducted in whole eye tissues samples from *Gclc* WT, *Gclc* HET, and *Gclc* KO mice at P3 and P21 days of age. (**A**) Hematoxylin and eosin-stained tissue sections used for IMS (Magnification = 100x). (**B**) Representative images indicating the intensity and localization of proteins. (**C**-**D**) Representative images indicating the intensity and localization of lipids in positive (**C**) and negative (**D**) mode. Individual images represent selected ions (m/z) and color intensities indicate relative abundances based on the provided color scale.

Analyses of the signal intensity of lipids within eyes of P3 day old mice revealed many differences between *Gclc* KO and *Gclc* WT eyes (Figs. 5C,D). Representative images taken in positive ion mode showed a decrease in the signal intensity of an unidentified lipid at m/z 256.8 and an increase in the signal intensity of an unidentified lipid at m/z 525.3 in the cornea of Gclc KO mice (Fig. 5C). A decrease in the distribution of putatively-identified phosphatidycholine (PC) (36:3) (m/z at 806.5) in the retina of *Gclc* KO mice was also observed (Fig. 5C). Analysis of the eyes in negative ion mode identified changes in several lipid species (Fig. 5D). Putatively-identified phosphatidylethanolamine (PE) (40:6) (m/z at 790.5) had decreased signal intensity in the retinas of *Gclc* KO and *Gclc* HET mice when compared with *Gclc* WT mice, however, *Gclc* KO mice have increased signal intensity when compared with *Gclc* HET mice. Conversely, the signal intensities for two putatively-identified phosphatidylinositol (PI) lipids (at m/z 863.6 and 885.6) increased in the cornea and retina of mice in which *Gclc* was deleted. Lastly, the signal intensity for putatively-identified NeuAcα2-3Galβ-Cer was increased in the retinas of *Gclc* KO mice relative to *Gclc* WT and *Gclc* HET mice (Fig. 5D).

### A conserved role for *gclc* in eye development

To further investigate the role of GSH in the developing eye, we developed *gclc* mutant knock-down (KD) and knock out (KO) zebrafish by using splice-blocking morpholinos (KD) or CRISPR/Cas9 genome editing (KO) in zebrafish embryos. Splice-blocking morpholinos (sbMOs) were developed that targeted either zebrafish *gclc* exon 1 (sb1) or zebrafish exon 5 (sb2) (Suppl. Fig. 1A). Administration of sb1 (3ng was determined to be the optimal dose) caused exon skipping and administration of sb2 (9ng was determined to be the optimal dose) caused intron retention (219 bp); these caused >90% and >60% reductions in *gclc* expression, respectively (Suppl. Fig. 1B). Administration of sb1 and sb2 resulted in a reduction in eye area by 26.5% and 10%, respectively (Fig. 6A) Representative brightfield images of *gclc* knock-down fish (3.5 days post fertilization (dpf)) after treatment with either sb1 or sb2 show the reduced eye area (Fig. 6A’). To genetically ablate *gclc*, CRISPR/Cas9 was utilized with a guide (g)RNA targeting *gclc* exon 3 (sgRNA1) (Suppl. Fig. 1A). Analyzing the resulting F0 *gclc* mutants, which are ≈ 30% mosaic mutants as indicated by CRISPR efficiency (Suppl. Fig. 1C), revealed that the eye area of 3.5 dpf *gclc* KO zebrafish is smaller than that of control zebrafish (Fig. 6B). Representative brightfield images of F0 *gclc* KO fish demonstrates the reduced eye size (Fig. 6B’).

**Figure 6.**
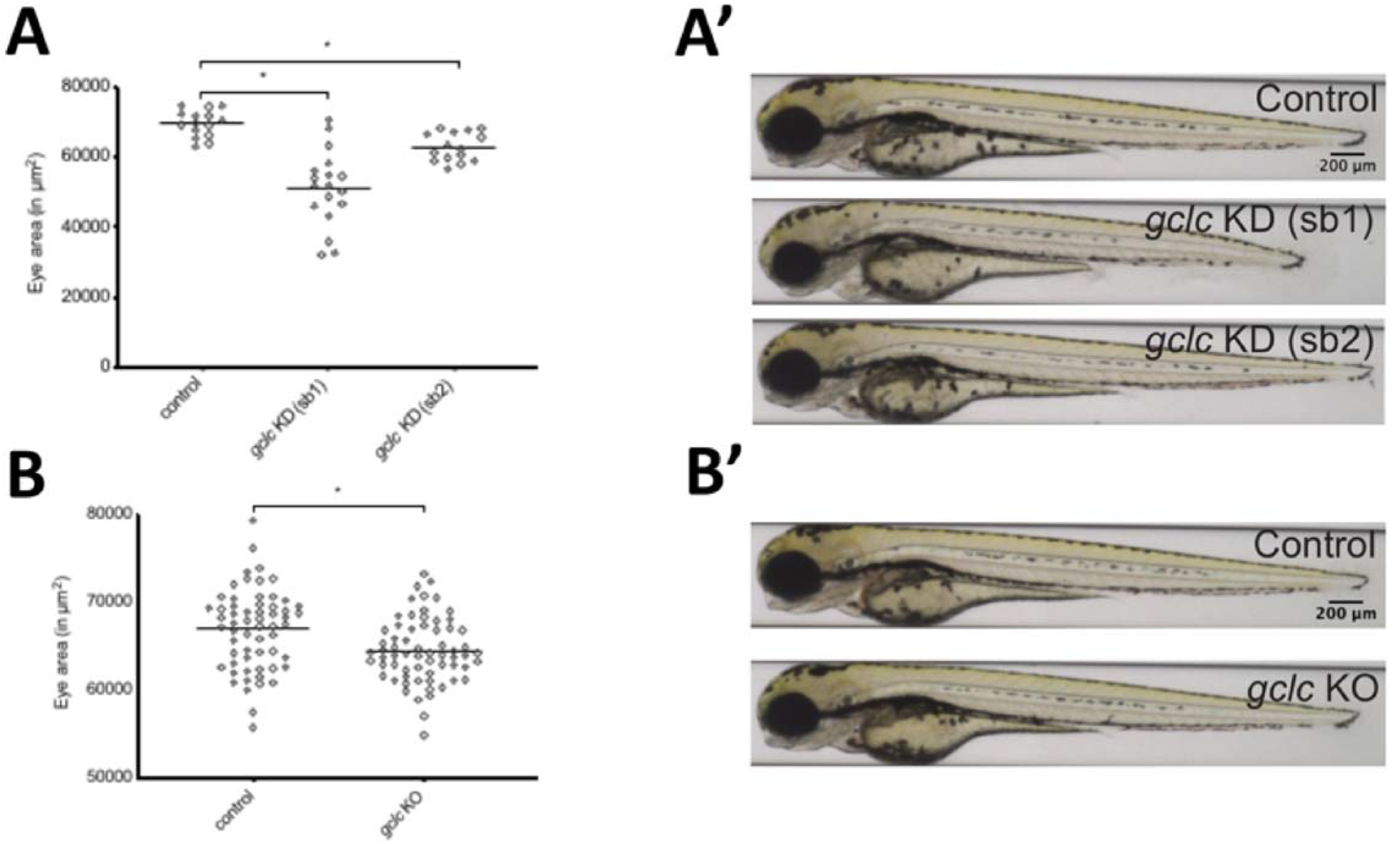
*gclc* gene suppression induces microphthalmia in zebrafish. (**A**) Embryonic eye area (in µm^2^) at 3.5 days post fertilization (dpf) of control and *gclc* KD fish, i.e., zebrafish in which *gclc* had been suppressed by sbMOs (*gclc* KD sb1 and *gclc* KD sb2). (**A’**) Representative brightfield image of the lateral view of 3.5 dpf control and *gclc* KD fish. (**B**) Embryonic eye area (in µm^2^) of 3.5 dpf control and *gclc* KO fish, i.e., zebrafish in which *gclc* had been suppressed using CRISPR (sgRNA1+Cas9). (**B’**) Representative brightfield image of the lateral view of 3.5 dpf control and *gclc* KO (sgRNA1+Cas9) fish. Each point in **A** and **B** represents results from an individual zebrafish. Horizontal lines indicate the mean. * P<0.05, one-way ANOVA with Tukey correction, compared to other group indicated.

## Discussion

In this study, we describe the creation and initial characterization of a conditional *Gclc* knock-out mouse model involving the use of the *Le-Cre* transgene (which results in gene deletion being limited to only the lens, cornea, conjunctiva, and eyelid). Additionally, we have described the creation of three *gclc*-depleted zebrafish models. GCLC is the rate limiting enzyme in glutathione (GSH) biosynthesis. As such, we expected *Gclc*-depleted animals would have reduced GSH abundance [24–26]. This expectation was confirmed in our *Gclc* KO mice. Thus, we demonstrate the creation of novel mouse and zebrafish models for the study of the impact of GSH depletion on eye development. Our results indicate that GSH is still present within the developing eye, but at a reduced amount. A failure to see a complete loss of GSH content in the lens was also observed in the LEGSKO mouse model [26]. Further, a large amount of GSH is produced in other ocular tissues besides the lens [40]. In summary, we have successfully deleted *Gclc*, GCLC protein expression is lost in surface ectoderm-derived tissues, and subsequently whole eye GSH content is reduced.

The microphthalmia phenotype occurring in our *Gclc* KO mice is starkly different from the cataract phenotype observed in another *Gclc* knock-out mouse model, the LEGSKO mouse model [26]. LEGSKO mice display reduced lenticular GSH content, elevated ROS, and cataract formation beginning as early as 4 months, but do not have microphthalmia. Analysis of the LEGSKO lens transcriptome revealed that GSH deficiency induces expression of detoxifying genes and activation of epithelial-mesenchymal transition (EMT) signaling [41]. Proteomic analysis of LEGSKO lenses further confirmed the activation of EMT signaling, changes in stress response proteins, and revealed a loss of lens specific markers [42]. The differences in phenotypes between these two mouse models may be explained by the fact that LEGSKO mice and *Gclc* KO mice have *Gclc* deletion at different developmental times, however, it is important to note that we did not analyze *Gclc* KO mice for cataract development at 4 months old. The LEGSKO mouse model relies on the *MLR10 Cre* transgene to drive gene deletion in lens-specific epithelial and fiber cells at approximately E10.5 [26, 43]. In contrast, our *Gclc* KO mouse model employs the *Le-Cre* transgene to drive gene deletion in surface ectoderm-derived tissues at approximately E9 [28]. Hence, it is conceivable that the timing of *Gclc* deletion, and subsequent resultant GSH depletion, may be vitally important for disrupting normal eye development.

Microphthalmia can be caused by developmental defects in both the anterior and posterior segments (total microphthalmia), or by defects in either the anterior or posterior segments (partial microphthalmia) [5]. *Gclc* KO mice display total microphthalmia, manifesting as severe developmental defects in both the anterior and posterior segments. The deformities observed in the posterior segment were unexpected because *Gclc* deletion was limited to only surface ectoderm-derived tissues. Mouse models of impaired lens development can display retinal malformations (i.e., infolding and expansion) [28, 45, 46]. Therefore, it is possible that the loss of *Gclc* from surface ectoderm-derived tissues is disrupting the developmental crosstalk between the lens and the retina.

*Gclc* KO eyes displayed an age-dependent phenotype that is observable from as early as E14.5 with a decreased anterior chamber size and retinal cellular layer expansion. By the time of weaning (P21), *Gclc* KO eyes have small vacuolated lens, severe retinal cellular layer expansion and in folding, hypoplasia of the iris, and corneal epithelial cell layers that are hyperproliferated and poorly differentiated. However, mice that have at least one *Le-Cre* transgene allele may develop microphthalmia regardless of any floxed alleles [36–39]. For example, mice homozygous for the *Le-Cre* transgene display high penetrance microphthalmia in a strain-dependent manner [38], whereas mice hemizygous for *Le-Cre* transgene display much lower penetrance in a similarly strain-dependent manner [36, 37]. FVB/N inbred mice served as the original background for the *Le-Cre* transgene [28] and *Le-Cre* hemizygous transgenic mice maintained on this background exhibit normal eye development. In contrast, a microphthalmia phenotype appears in *Le-Cre* hemizygous transgenic mice as the genetic contribution of FVB/N is reduced [37, 38]. Given that the *Le-Cre* transgene utilizes the *Pax P0* promoter, others have postulated that it may deplete cofactors required for endogenous *Pax6* expression [37]. However, a recent study was unable to confirm this hypothesis [36]. Instead, it was found that the microphthalmia phenotype (in *Le-Cre* homozygous mice) was caused by changes in the expression of genes involved in the negative regulation of cell proliferation and cell growth, and positive regulation of apoptosis pathways. Since these two studies used mice of different strains, the disagreement between them may be explained by strain-dependent differences in *Pax6* expression [48] and/or susceptibility to eye development phenotypes when exposed to changes in *Pax6* expression or PAX6 levels [48, 49]. Regardless of its cause, it is critical that any study utilizing the *Le-Cre* transgene control for the effect of the *Le-Cre* alone. Isolating the *Le-Cre* transgene on the mixed B6/FVB background (as used throughout this study) confirmed that the *Le-Cre* transgene alone causes microphthalmia (Fig. 3B). The present study demonstrates that *Gclc* deletion during early eye development exacerbates the microphthalmia phenotype of the *Le-Cre* transgene alone. Our additional observations that *Gclc* HET mice display unilateral microphthalmia and *gclc* deletion in zebrafish causes microphthalmia underscore an important role for *Gclc* expression in eye development.

Preliminary molecular investigations in *Gclc* KO eyes revealed a decrease in a cell proliferation marker and increased apoptosis in the developing lens. Cells must proliferate and go through apoptosis for proper organogenesis [50]. Many studies indicate that lens development requires proper cellular proliferation [51–53] and apoptosis [51, 54]. The lens epithelium produces precursor lens fiber cells [55], therefore, a decrease in proliferation of lens epithelial cells will result in a reduction of the lens fiber mass, as observed in *Gclc* KO mice. Further contributing to the decreased lens fiber mass was an increase in apoptosis in lens epithelial cells and secondary fiber cells. Other studies have shown that changes in PAX6 [56] and β-catenin [57] expression may cause a loss of lens epithelium proliferation and increased cell death. However, we did not observe changes in the expression of either of these two proteins. A change in the translation of αA-crystallin splice variants in the lenses of *Gclc* KO mice at P0 was observed. The significance of this within the context of microphthalmia, and the cause of increased cell death and decreased proliferation all remain to be elucidated.

Although the overall phenotype observed in *Gclc* KO mice differ from those of the LEGSKO mouse model [26, 41, 42], they are similar to other animal models of oxidative stress. Hydrogen peroxide treatment of lenses obtained from adult glutathione peroxidase (GSHPx) knock-out mice display vacuolated lens epithelial cells in the central region, and degenerated nuclei, organelle loss, and elevated DNA strand breaks in the equatorial region of the lens epithelial cells [58]. Similarly, genetic suppression of glutaredoxin 2 (GLRX2) expression (i.e., knock-out) increased the sensitivity of ex vivo mouse lens epithelial cell to oxidative stress and reduced their viability as a result of increased glutathionylation [59]. It is possible that such a process may be occurring within the *Gclc* KO mice to produce the observed microphthalmia phenotype.

Imaging mass spectrometry (IMS) allows the spatial characterization of thousands of proteins and other small molecules in a tissue *in situ* during a single experiment [60] (for review, see [61]). This technique has been utilized in investigations of the cornea [62], retinal lipids [63, 64], and lens GSH [65, 66], lipids [67–69], UV-filters [70], and proteins [71, 72]. To our knowledge, our study is the first to use IMS in a microphthalmia mouse model. IMS of eye tissue revealed changes in the distribution and intensities of thymosin β4, cytochrome C oxidase 7C, and histone H2B. Thymosin β4 has a regenerative role (by down-regulating inflammation, inducing tissue regeneration and inhibiting scarring) in the skin, heart, brain and eye [73]. In the eye, thymosin β4 promotes corneal wound healing by decreasing inflammation [74], regulating nuclear factor-kappa B (NF-κB) in the immune response [75] and regulating matrix metalloproteases [76]. Therefore, the observed loss of thymosin β4 expression from the cornea of *Gclc* KO mice may render the eyes of these mice with a decreased capacity to respond to exogenous stressors and to heal. Cytochrome C oxidase 7C (Cox7C), a nuclear encoded subunit of the COX holoenzyme, is highly abundant in mouse retinas [77–79]. It is the terminal enzyme of the electron transport system [80]. Histone H2B is one subunit of the histone H2A-H2B heterodimer [81], a key histone component [82]. Histone H2B is up-regulated in zebrafish retina undergoing regeneration [83]. These proteins are highlighted as markers of the retina, which clearly demonstrate the retinal expansion observed by H&E staining. IMS analyses also revealed marked changes in the ocular lipid profiles between *Gclc* KO mice. Phosphatidylinositols (PIs) regulate many processes ranging from; cellular proliferation to death [84]. PIs play a role in lens structural intengrity and photoreceptor cell survival in zebrafish [85]. The retinal expansion may explain the increased signal intensity for PI(38:4) within the retina of *Gclc* KO mice; perhaps they are also important for cell survival within the outer nuclear, inner nuclear, inner plexiform, and ganglion cell layers. It is possible that if *Gclc* KO mice were observed for long enough that they too would develop a macular dystrophy phenotype. The role of PI(36:1) in corneal tissues is unknown, however, it may also regulate cell signaling and could explain the corneal abnormalities observed. While, PIs signal intensities did not differ in the lenses of *Gclc* KO mice, phosphatidylinositol 3-kinase is necessary for the survival and differentiation of lens fiber cells [86]. Phosphatidylcholines (PCs) are important for rod function, longevity, and function; a shift in their composition is associated with a mouse model for Stargardt-like macular dystrophy [87]. Gangliosides are also known to be abundant throughout the retina, and change as the retina matures [88]. The increased NeuAcα2-3Galβ-Cer signal intensity in *Gclc* KO retinas may result from the retinal layer expansion. It is hoped that with continued studies of lipids throughout eye development by IMS, that the role for *Gclc* in regulating ocular lipid composition will be elucidated.

*Gclc* is highly expressed in the zebrafish eye throughout embryonic development [29], and is involved in the zebrafish antioxidant response to xenobiotic exposure [89–91]. Herein, we present a novel role for *gclc* in regulating zebrafish eye development; a reduction in *gclc* expression causes reduced eye area (microphthalmia). Thus, demonstrating a role for *Gclc* in regulating eye development that is conserved throughout the chordate family. The fact that *gclc* KD fish display a more severe phenotype than *gclc* KO fish can be explained by the mosaicism of *gclc* KO fish and the genetic compensation mechanisms activated by gene KO systems but not gene KD systems [92, 93]. Our novel *gclc* deficient zebrafish models may be used to provide important insights into human eye development, similar to the recently-described *dnajb1a* zebrafish model. In patients with Peter’s Anomaly, DNAJB1 transcripts were shown to be a target of FOXE3. Zebrafish deficient in *dnajb1a* (the zebrafish *DNAJB1* ortholog) were used to confirm the role of DNAJB1 in this condition [94]. It is anticipated that *gclc-*depleted zebrafish will similarly provide unique insights into human lens development and thereby provide benefits for human health.

The present study provides compelling evidence that the loss of *Gclc* results in severe microphthalmia which is associated with profound molecular changes. In so doing, it represents the first description of a novel role for *Gclc* in eye development. Thus, *Gclc* KO mice and *gclc-*depleted zebrafish are valuable new models for studying microphthalmia. Investigations in these animal models, utilizing powerful next-generation technologies (IMS, RNA-seq, and redox proteomics), are ongoing. It is expected that these studies will accelerate the development of new strategies for the prevention of and treatment of microphthalmia.

