## Supplemental Figure 1 for "*Gclc* deletion in surface-ectoderm tissues induces microphthalmia"

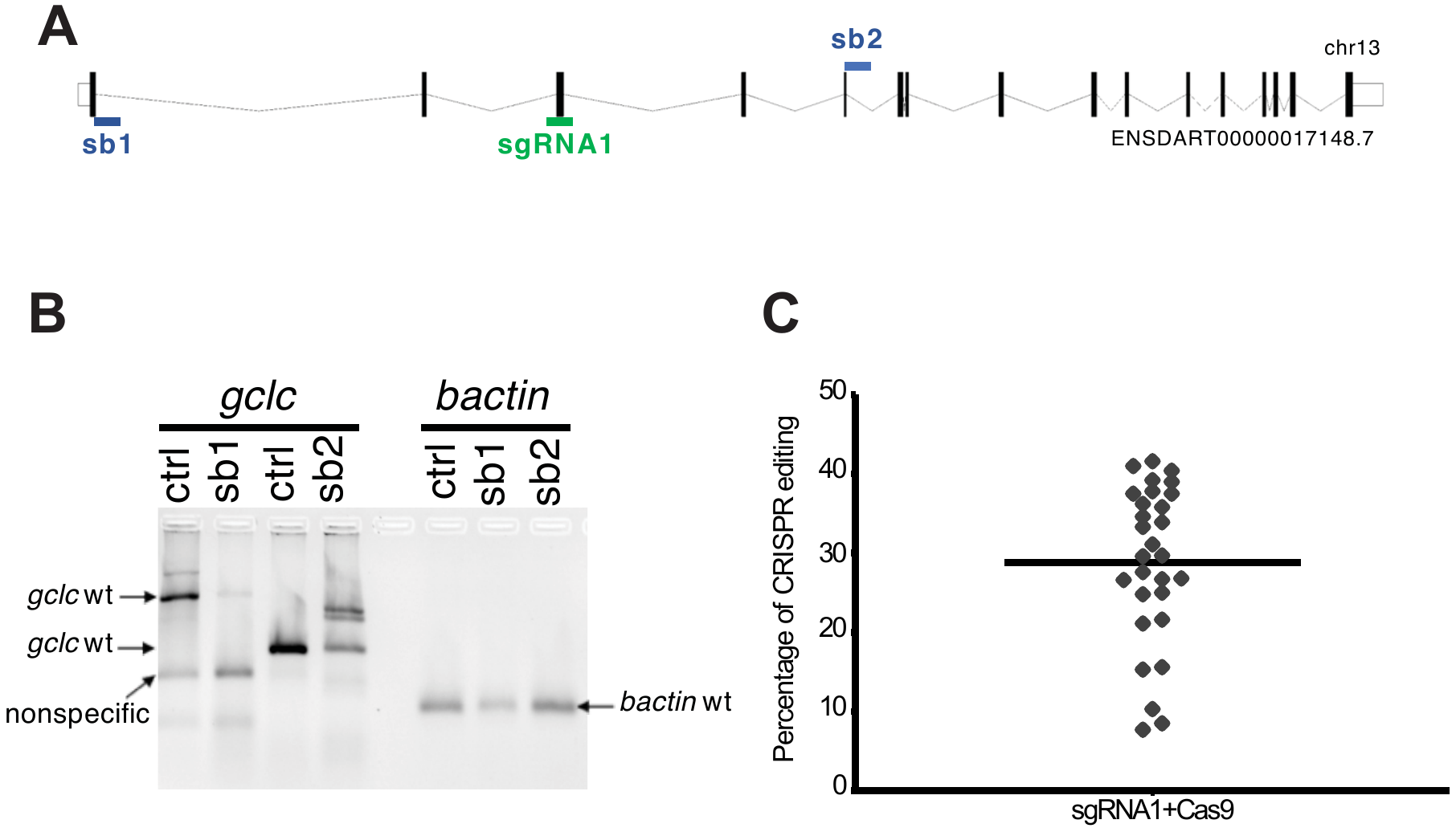

**Figure S1: *gclc* targeting details and efficiencies**

(**A**) Schematic of the *Danio rerio* *gclc* locus with coding exons (block boxes), introns (lines), and untranslated regions (white boxes). The position of the CRISPR (sgRNA1) target is indicated by a green line. NSDART00000017148.7 refers to the transcriptID in Ensembl. (**B**) RT-PCR of *gclc* and *bactin* at 3 dpf from whole body of control (ctrl) and *gclc-*suppressed (sb1 and sb2) zebrafish. The two separate bands for *gclc* WT result from the use of two different primer sets. (**C**) Crispr efficiency (in percentage) from *gclc* KO (sgRNA1+Cas9) treated zebrafish.
